# Epigenomic programming of peripheral monocytes determines their transcriptional response to the tumor microenvironment

**DOI:** 10.1101/2024.06.19.599675

**Authors:** Máté Kiss, Laszlo Halasz, Eva Hadadi, Wilhelm K. Berger, Petros Tzerpos, Szilard Poliska, Daliya Kancheva, Ayla Debraekeleer, Jan Brughmans, Yvon Elkrim, Liesbet Martens, Yvan Saeys, Bence Daniel, Zsolt Czimmerer, Damya Laoui, Laszlo Nagy, Jo A. Van Ginderachter

## Abstract

Classical monocytes are recruited to tumors and undergo transcriptional reprogramming resulting in tumor-promoting functions. Epigenomic features, such as post-translational modification of histones and chromatin accessibility, are key determinants of transcription factor binding and thereby play an important role in determining transcriptional responses to the tissue environment. It is unknown to what extent the epigenetic landscape of peripheral monocytes is rewired by cancer and how this could shape their transcriptional response upon recruitment to the tumor microenvironment. Here we used a combination of genome-wide assays for mRNA expression, chromatin accessibility and multiple histone modifications (H3K4me1, H3K4me3, H3K27ac) in a mouse model to investigate changes in the epigenomic landscape of peripheral monocytes. We then linked these epigenetic alterations to gene expression changes in monocytes occurring in the periphery or during tumor infiltration. We found that the distal tumor caused extensive remodeling of both H3K4me3^+^ promoters and H3K4me1^+^ enhancers in peripheral monocytes. Specifically, this involved the repression of interferon-responsive promoters and enhancers as well as the establishment of enhancers harboring binding motifs for transcription factors downstream of inflammatory and cytokine signaling pathways. The enhancers altered in the periphery could be linked to sustained gene expression changes which were less likely to be reversed in the tumor microenvironment. In addition, genes activated upon tumor infiltration showed prior epigenetic priming in peripheral monocytes. Overall, these results indicate that the epigenomic landscape of peripheral monocytes is altered in response to a distal tumor, and this could shape the transcriptional response of monocytes when they encounter microenvironmental signals upon infiltrating the tumor.

## Introduction

Classical monocytes play an essential role in host defense through their ability to rapidly migrate to sites of injury or infection where they initiate an inflammatory response (1). Similar to inflamed tissues, classical monocytes are mobilized to tumors where they either differentiate into tumor-associated macrophages or accumulate without differentiation, depending on the cytokine milieu (2, 3). Both monocytes and monocyte-derived macrophages can exert tumor-promoting activities, including suppression of anti-tumor T-cell responses, promotion of angiogenesis and facilitation of metastatic colonization (3). These activities arise due to extensive transcriptional reprogramming of monocytes when transitioning from the blood to the tumor, involving the activation of genes encoding inflammatory cytokines, pro-angiogenic factors and growth factors (4–7). However, recruited monocytes can also acquire an immunostimulatory phenotype and promote anti-tumor immunity in certain contexts. Specifically, the presence of type I interferon in the tumor can induce an immunostimulatory phenotype which supports the antitumor T-cell response (8, 9). Such interferon-responsive monocytes can also be detected during the course of successful cancer immunotherapies and their abundance is associated with a better response to treatment (9–12). Overall, these observations indicate that the transcriptional response of monocytes to microenvironmental signals is paramount in determining their phenotype and consequently their impact on disease outcome.

It is increasingly appreciated that cancer can disrupt immune homeostasis not only locally but also systemically (13). Indeed, multiple studies have demonstrated that modulation of monocyte function in cancer occurs already prior to their infiltration into the tumor (14). Phenotypic changes that have been observed in multiple tumor types include the acquisition of immunosuppressive activity and reduced responsiveness to interferon stimulation (14–18). Peripheral blood monocytes from cancer patients show a distinct gene expression profile compared to healthy controls and these changes are largely cancer type-specific (19, 20). Importantly, transcriptional alterations in blood monocytes can also be detected in patients with localized early-stage tumors (20).

Although transcriptomic changes in peripheral monocytes in the context of cancer are well documented, it is currently unknown to what extent the epigenomic landscape of monocytes is altered by a distant tumor. Epigenomic features, such as post-translational modification of histones and chromatin accessibility, are key determinants of DNA binding and thus genome-wide localization and activity of transcription factors activated by external signals (21). The preformed epigenomic landscape determined by these features plays a key role in directing transcriptional responses to the cues of the microenvironment. Importantly, epigenetic marks are typically less dynamic than the activity of upstream signaling pathways and can therefore convert transient signals into long-lasting transcriptional alterations or transcriptional memory by acting as bookmarks which could persist even after cessation of the stimulus (22, 23). The pre-formed epigenomic landscape is likely to be important in shaping the responses of monocytes when they encounter new signals upon tumor infiltration. Nonetheless, it remains unknown whether some tumor-induced transcriptional changes already in peripheral monocytes could be programmed at the epigenomic level and whether epigenomic changes could prime genes for activation upon tumor infiltration.

In this study, we aimed to address these knowledge gaps in a murine model by using an array of complementary genome-wide assays for mRNA expression (RNA-seq), histone modifications (H3K4me3 CUT&Run, H3K4me1 and H3K27ac CUT&Tag) and chromatin accessibility (ATAC-seq) to detect potential cancer-induced changes in the epigenomic landscape of peripheral monocytes and link these to gene expression changes occurring in the periphery or upon tumor infiltration.

## Results

### Tumor-induced transcriptional alterations in peripheral monocytes persist following tumor infiltration and involve repression of interferon-response genes

In order to determine potential links between peripheral alterations in monocytes and their local reprogramming in the tumor, we sought to identify genes whose expression was modulated by the tumor either in the periphery or upon tumor infiltration. To this end, we performed bulk RNA-sequencing on CD11b^hi^ Ly6C^hi^ Ly6G^-^ MHC-II^-^ classical monocytes isolated from the tumor and blood of Lewis lung carcinoma (LLC) tumor-bearing mice as well as from the blood of healthy syngeneic mice (**Figure 1A**). The comparison between cells from genetically identical tumor-bearing and healthy mice allowed us to exclude the impact of genetic variation and precisely identify tumor-induced changes in the gene expression of peripheral monocytes (**Figure 1A**). The LLC model was chosen as this tumor recruits a large number of monocytes which show immunosuppressive features (24).

**Figure 1.**
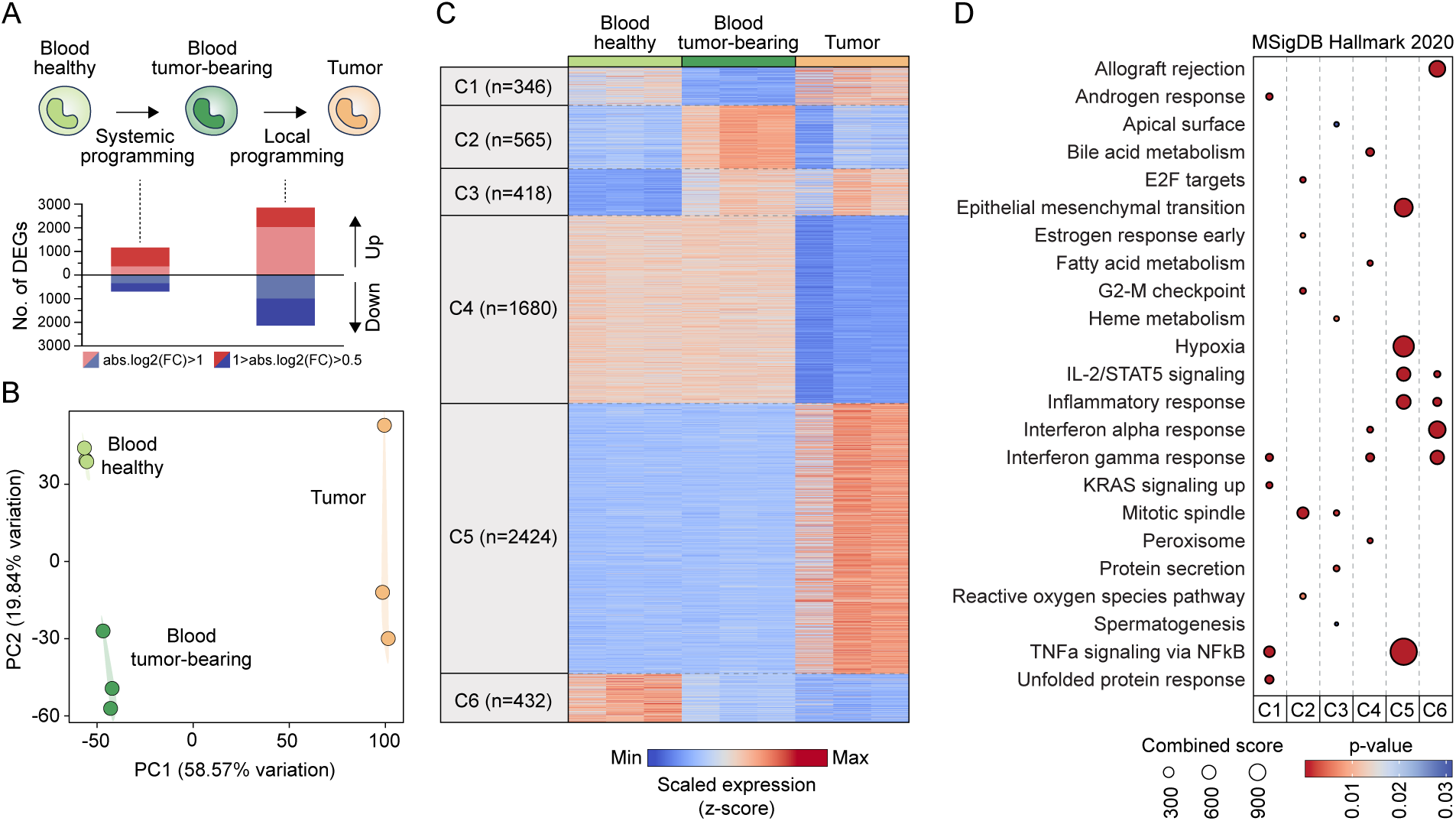
Tumor-induced transcriptional alterations in peripheral monocytes persist following tumor infiltration and involve repression of interferon-response genes. (A) Experimental scheme to examine peripheral and local programming of monocytes by the tumor and bar graphs showing the number of differentially expressed genes (DEGs) for the indicated comparisons, with different absolute log2(fold change) value thresholds. (B) Principal-component analysis of RNA-seq profiles of Ly6C^hi^ monocytes sorted from the blood of healthy mice, blood of LLC tumor-bearing mice and LLC tumors (n=3). (C) Heatmap showing genes that were differentially expressed in at least one comparison between different conditions (P<0.05 and log2[fold change]≥0.5), and separated to six clusters by k-means clustering. (D) Enrichment of MSigDB Hallmark gene sets in the different gene expression clusters. Top 5 enriched gene sets are shown for each cluster based on combined score.

We found that the transcriptome of tumor-infiltrating monocytes was very different from monocytes in the blood, while the tumor also induced distinct, albeit less pronounced, changes in the transcriptome of peripheral monocytes (**Figure 1A,B**). Differentially expressed genes (P<0.05 and log2[fold change]≥0.5), could be split into six major clusters with distinct gene expression patterns (**Figure 1C, Table S1**). This approach enabled us to identify two gene clusters (Cluster 3 and 6) whose expression was modulated in the periphery in tumor-bearing mice and these alterations persisted following tumor infiltration. Cluster 3 included genes whose expression was induced by the tumor in peripheral monocytes and their expression level remained elevated following tumor infiltration. Similarly, Cluster 6 included genes whose expression was repressed in blood monocytes of tumor-bearing mice and remained repressed following tumor infiltration. We also identified gene clusters (Cluster 1 and 2) whose expression was altered in peripheral monocytes in response to the distal tumor, but this was reversed upon tumor infiltration. Finally, we found two large gene clusters (Cluster 4 and 5) whose expression did not show major differences in the blood, but showed marked down- or upregulation during blood-to-tumor transition.

In order to link functional gene programs to these gene expression clusters, we analyzed the enrichment of Hallmark gene sets (**Figure 1D**). Genes induced in peripheral monocytes of tumor-bearing mice (Cluster 2 and 3) showed specific enrichment of proliferation-related gene sets ("G2-M checkpoint", "Mitotic spindle", "E2F targets"), suggesting enhanced monocyte production in response to the tumor, in line with previous observations (15). Cluster 3 genes, which were induced by the tumor in the periphery and remained induced within the tumor, also included genes related to protein secretion and hem metabolism, the latter being an important driver of macrophage reprogramming in tumors (25–27). Genes that were only turned on upon tumor infiltration (Cluster 5) showed specific enrichment for gene sets related to inflammation and the hypoxia response, indicating local activation and adaptation to the tumor microenvironment. We found that the gene cluster repressed upon tumor-infiltration (Cluster 4) was enriched for interferon response genes. Interestingly, however, interferon response-related genes showed the strongest enrichment in the gene cluster which was already repressed in the periphery and remained repressed following tumor infiltration (Cluster 6).

Overall, these results demonstrate that some gene expression changes in peripheral monocytes induced by a distal tumor persist following tumor infiltration, suggesting a tumor-driven preconditioning of monocytes impacting their transcriptional program within the tumor.

### Tumor-induced epigenetic alterations at promoters of peripheral monocytes lead to repression of interferon-responsive promoters and activation of key pro-tumoral genes

Our observation that some peripheral blood monocyte genes showed durable expression changes in the tumor-bearing state suggested that some of these genes were programmed at the epigenetic level in the periphery, making their transcriptional induction or repression more stable in the tumor.

In order to detect potential tumor-induced epigenomic alterations in peripheral monocytes, we first assessed the genome-wide distribution of Histone H3 Lysine 4 trimethylation (H3K4me3), a key histone modification for transcriptional initiation characteristic of promoter regions (28), using the CUT&Run method. We detected H3K4me3 signal predominantly (88%) at or near promoter regions, as expected (**Figure S1A**). Comparative analysis between blood monocytes from LLC tumor-bearing and healthy mice revealed a set of promoters with altered H3K4me3 signal (166 induced and 312 repressed sites with P<0.05, log2[fold change]≥0.5) (**Figure 2A, B, Table S2**).

**Figure 2.**
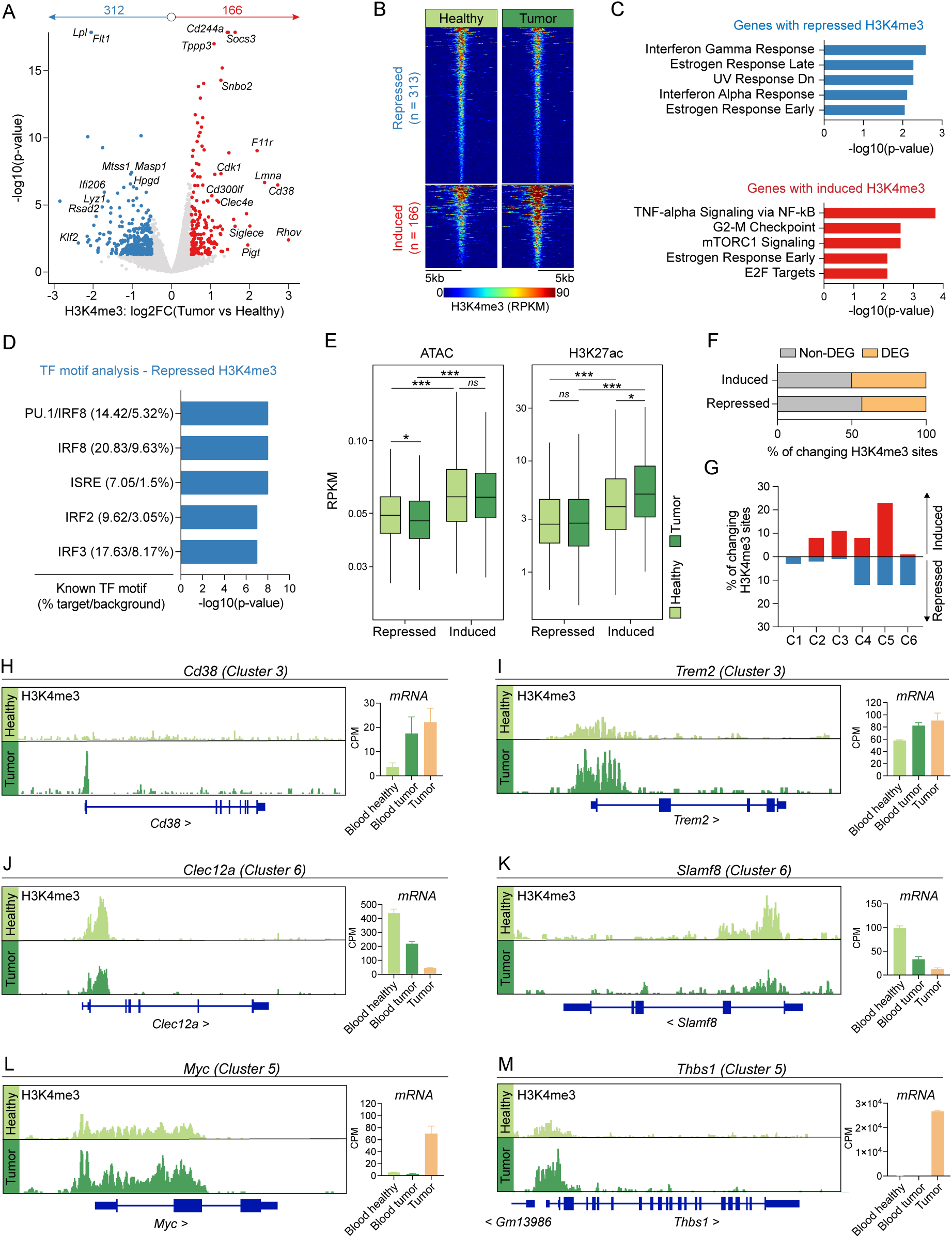
Tumor-induced epigenetic alterations at promoters of peripheral monocytes lead to repression of interferon-responsive promoters and activation of key pro-tumoral genes. (A) Volcano plot showing genomic sites with differential H3K4me3 signal between Ly6C^hi^ monocytes from the blood of LLC tumor-bearing versus healthy mice (P<0.05 and log2[fold change]≥0.5). Representative closest protein-coding genes are highlighted. (B) Read distribution plot showing H3K4me3 signal at genomic sites with differential H3K4me3 signal. (C) Enrichment of MSigDB Hallmark gene sets among genes annotated to genomic sites with differential H3K4me3 signal. Top 5 gene sets are shown by p-value. (D) Top 5 known transcription factor (TF) motifs enriched among genomic sites showing repressed H3K4me3 signal in Ly6C^hi^ monocytes from the blood of LLC tumor-bearing mice versus healthy mice. (E) Chromatin accessibility (ATAC-seq) and H3K27ac signal in Ly6C^hi^ monocytes from the blood of healthy and LLC tumor-bearing mice at genomic sites with repressed or induced H3K4me3 signal when comparing tumor-bearing vs. healthy. Boxes show the median and the interquartile range (IQR), whiskers show highest and lowest values within 1.5xIQR. Wilcoxon test; ns: not significant, *P<0.05, **P<0.01, ***P<0.001. (F) Frequency of genes with differential H3K4me3 signal that are among the differentially expressed genes identified in Figure 1C (DEG) or not (non-DEG). (G) Frequency of different cluster identities (determined in Figure 1C) among the genes with induced or repressed H3K4me3 signal. (H-M) Genome browser snapshots showing average H3K4me3 signal in Ly6C^hi^ monocytes from the blood of healthy and LLC tumor-bearing mice at representative genes from different expression clusters. Bar plots show mRNA expression of the same genes determined by RNA-seq. Bar plots show mean+SEM.

By linking the nearest gene to each changing promoter, we found that genes with induced promoters were related to cellular functions such as cell division (e.g. *Cdk1*, *Cdkn1a*, *Plk1*), cell adhesion and migration (e.g. *Rhov*, *F11r*), and immunosuppression (e.g. *Socs3*, *Siglece*, *Cd38*). Among the genes with repressed promoters, we found several genes linked to the interferon response (e.g. *Ifit206*, *Rsad2*, *Mx2*) (**Figure 2A**). These observations were further supported by the significant enrichment of the corresponding Hallmark gene sets related to cell division and the interferon response among the genes with induced and repressed promoters, respectively (**Figure 2C**).

While no known motifs were significantly enriched among induced promoters, we found significant enrichment of interferon-responsive IRF and ISRE motifs among repressed promoters (**Figure 2D**). In line with these data, nearly all IRF transcription family members, except for *Irf9,* showed reduced expression in peripheral monocytes of tumor-bearing mice, with *Irf1* and *Irf7* showing the greatest downregulation (**Figure S1B**).

We then performed ATAC-seq to map chromatin accessibility across the genome of blood monocytes from healthy and LLC tumor-bearing mice and used these data to compare chromatin accessibility specifically at differential promoters. This analysis showed that average chromatin accessibility slightly decreased at repressed promoters in tumor-bearing mice (**Figure 2E**). In addition, promoters that were induced by the tumor showed higher chromatin accessibility already at baseline in healthy mice and accessibility at these sites did not increase further in response to the tumor (**Figure 2E**).

In order to obtain information about promoter activity, we used CUT&Tag to analyze the genome-wide distribution of Histone H3 Lysine 27 acetylation (H3K27ac), a histone modification characteristic of promoters and enhancers with active transcription (29–32). Similar to chromatin accessibility, promoters that were induced in tumor-bearing mice already showed higher average H3K27ac signal in healthy mice compared to repressed promoters, indicating higher baseline activity. In addition, the H3K27ac signal was further enhanced on this set of promoters in response to the distal tumor (**Figure 2E**).

Next, we asked whether genes with differential promoters showed changes in their expression either in the blood or upon tumor infiltration and, if so, which previously identified gene expression cluster (shown in Figure 1C) they can be linked to. Remarkably, less than half of the genes with altered promoters showed expression changes in the blood or upon tumor infiltration, indicating that promoter alterations were not necessarily associated with a direct change in transcription (**Figure 2F**). Among the genes where promoter reprogramming could be linked to transcriptional change, we found that genes induced in the circulation (Clusters 2 and 3 in Figure 1C) could be almost exclusively linked to induced promoters (**Figure 2G**). Such epigenetically programmed genes included *Cd38* and *Trem2* which have well-established roles in driving immunosuppressive and pro-tumoral functions of monocytes/macrophages (33–40). These genes showed increased H3K4me3 at their promoters in the circulating monocytes from tumor-bearing mice, and this was associated with a persistently enhanced transcription upon tumor infiltration (**Figure 2H,I**). Similarly, repressed genes in the circulation (Clusters 1 and 6 in Figure 1C) could only be linked to repressed promoters (**Figure 2G**), further indicating a close correspondence between the direction of changes in promoter H3K4me3 levels and in gene expression. Such repressed genes included *Slamf8*, a negative regulator of ROS production and *Clec12a*, an inhibitory receptor involved in cell death sensing (**Figure 2J,K**) (41, 42).

Interestingly, genes that were upregulated only upon tumor infiltration (Cluster 5 in Figure 1C) could be preferentially linked to promoters induced in the circulation (**Figure 2G**). This raised the possibility that tumor-induced promoter activation could prime genes for activation upon tumor infiltration. This gene set included *Myc* and *Thbs1*, which have been previously linked to tumor-promoting functions in macrophages (**Figure 2L,M**) (43–45).

Overall, these data indicate that a set of promoters exhibit tumor-induced alterations in monocytes, prior to tumor infiltration. This involves the repression of interferon-responsive promoters and the priming of key pro-tumoral genes for expression in the tumor microenvironment.

### The enhancer landscape in peripheral monocytes is remodelled in response to the tumor

We hypothesized that enhancers of peripheral monocytes may also show changes in response to the distal tumor. Hence, we used CUT&Tag to analyze the enhancer landscape of peripheral monocytes from healthy and LLC tumor-bearing mice by assaying the genome-wide distribution of histone H3 Lysine 4 monomethylation (H3K4me1), a histone modification which marks enhancers (46).

Differential analysis between peripheral monocytes from tumor-bearing and healthy mice revealed more than three thousand genomic regions with altered H3K4me1 signal, which represented approximately 10% of all detected H3K4me1^+^ enhancers (1757 induced and 1427 repressed sites at P<0.05, log2[fold change]≥0.5 among 33110 total detected H3K4me1^+^ regions) (**Figure 3A,B, Table S3**). Most of these tumor-modulated enhancers were located in distal intergenic regions or introns (**Figure 3C**).

**Figure 3.**
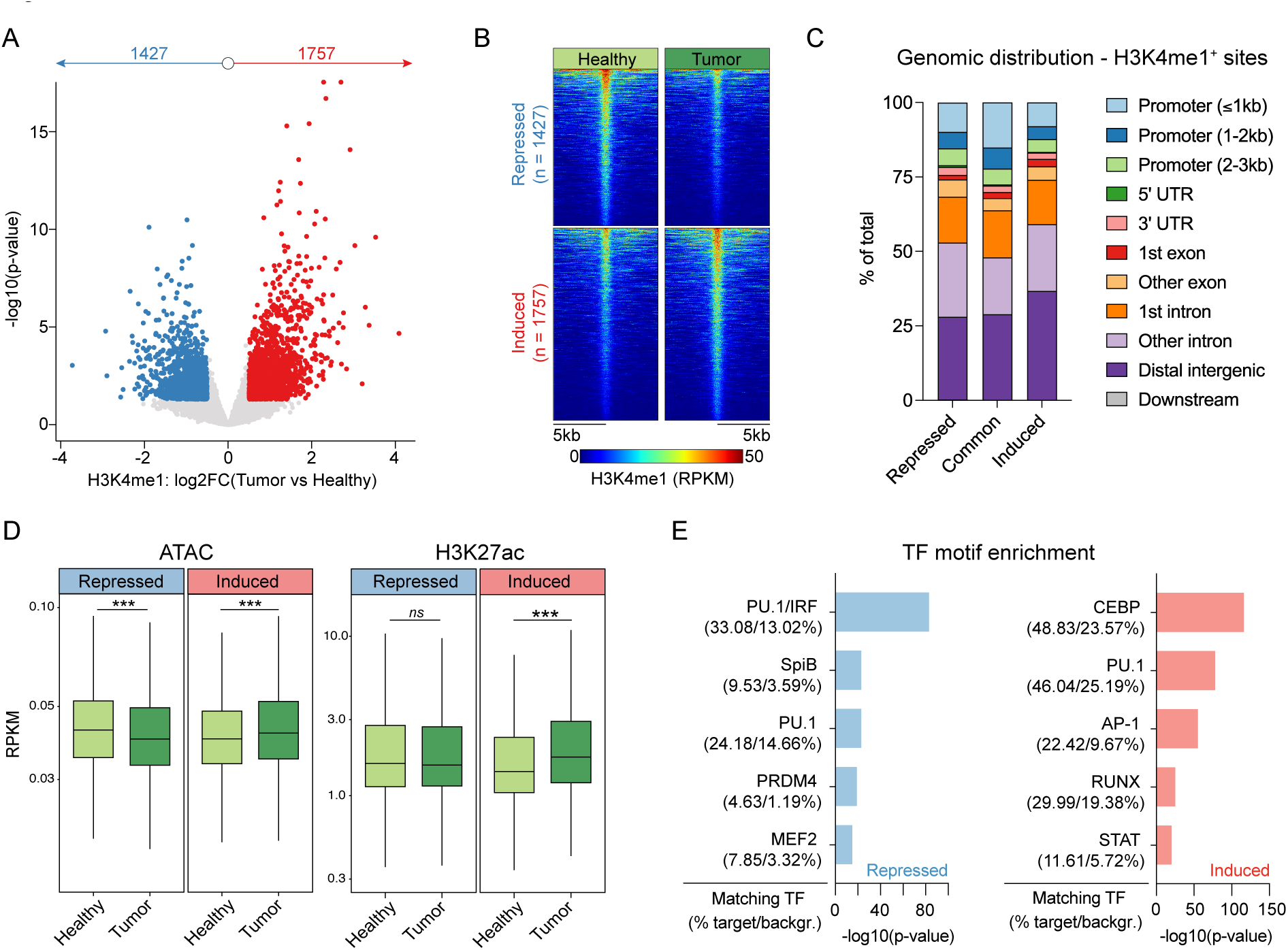
The enhancer landscape in peripheral monocytes is remodelled in response to the tumor. (A) Volcano plot showing genomic sites with differential H3K4me1 signal between Ly6C^hi^ monocytes from the blood of LLC tumor-bearing versus healthy mice (P<0.05 and log2[fold change]≥0.5). (B) Read distribution plot showing H3K4me1 signal at genomic sites with differential H3K4me1 signal. (C) Genomic distribution of sites with differential and unaltered H3K4me1 signal. (D) Chromatin accessibility (ATAC-seq) and H3K27ac signal in Ly6C^hi^ monocytes from the blood of healthy and LLC tumor-bearing mice at genomic sites with repressed or induced H3K4me1 signal when comparing tumor-bearing vs. healthy. Boxes show the median and the interquartile range (IQR), whiskers show highest and lowest values within 1.5xIQR. Wilcoxon test; ns: not significant, ***P<0.001. (E) Top 5 transcription factors (TF) matched to *de novo* motifs enriched among genomic sites showing repressed or induced H3K4me1 signal in Ly6C^hi^ monocytes from the blood of LLC tumor-bearing mice vs. healthy mice. Motifs are shown in Figure S2A.

Average chromatin accessibility significantly decreased at repressed enhancers, whereas it increased at induced enhancers, indicating chromatin remodeling at these sites (**Figure 3D**). Average H3K27ac levels significantly increased at induced enhancers in tumor-bearing mice, indicating elevated enhancer activity, whereas H3K27ac levels remained unaltered at repressed enhancers (**Figure 3D**).

Transcription factor motif enrichment analysis on differential enhancers revealed enrichment of the PU.1 motif both at repressed and induced sites (**Figure 3E, Figure S2A**). This was expected, as PU.1 is a myeloid lineage-determining factor which has an essential role in maintaining cell type-specific enhancers (21). Among the repressed enhancers, we found strong enrichment of the PU.1-IRF composite motif, adding further support to the notion that IRF activity is suppressed in peripheral monocytes in response to the tumor (**Figure 3E**). Among the induced enhancers, the most significantly enriched motifs could be linked to CEBP, AP-1, RUNX and STAT transcription factor families, all of which can mediate response to extracellular inflammatory signals and cytokines (**Figure 3E**). Expression levels of transcription factors belonging to these families were largely unaltered in peripheral monocytes, but several of them showed marked upregulation following tumor infiltration, such as *Cebpb*, *Jun*, *Junb*, *Fosb*, *Fosl2*, *Atf3* and *Runx3* (**Figure S2B**).

Altogether these results indicate that a distal tumor is able to remodel the enhancer landscape in peripheral monocytes. Specifically, this involves repression of enhancers capable of binding interferon-responsive IRF transcription factors as well as the establishment of enhancers harboring motifs for transcription factors that can be activated by inflammatory and cytokine signaling.

### Altered enhancers in peripheral monocytes are associated with distinct gene expression patterns upon tumor infiltration

Finally, we sought to link differential enhancers to genes modulated by the tumor either in the periphery or upon tumor infiltration. To this end, we analyzed the frequency of induced and repressed enhancers in a +/-100 kilobase window around the transcription start site of differentially expressed genes belonging to different clusters described in Figure 1C (**Figure 4A**). The observed enhancer frequencies were then compared to enhancer frequencies detected around random gene sets of the same size, in order to determine the enrichment or depletion of differential enhancers around distinct gene clusters (**Figure 4A**). This analysis revealed the strongest enrichment of induced and repressed enhancers around the previously defined C3 and C6 gene expression clusters, respectively (**Figure 4B, Figure S3**). These clusters comprised genes that showed induction or repression in the periphery, which persisted following tumor infiltration (e.g. *Atp11a* and *Man2a1* from C3; *Ifi27l2a* and *Tlr11* from C6, **Figure 4C-F**). Interestingly, genes whose peripheral activation or inhibition could be reversed in the tumor (C2 and C1, respectively) did not show such a strong association with altered enhancers (**Figure 4B, Figure S3**). Notably, genes that were only induced in the tumor microenvironment (C5) showed a marked enrichment of induced enhancers and a significant depletion of repressed enhancers in their vicinity (e.g. *Mapk6* and *Rhov*, **Figure 4B,G,H, Figure S3**). In addition, we also found significant enrichment of repressed enhancers but no depletion of induced enhancers, around genes which were downregulated upon tumor infiltration (C4) (**Figure 4B, Figure S3**).

**Figure 4.**
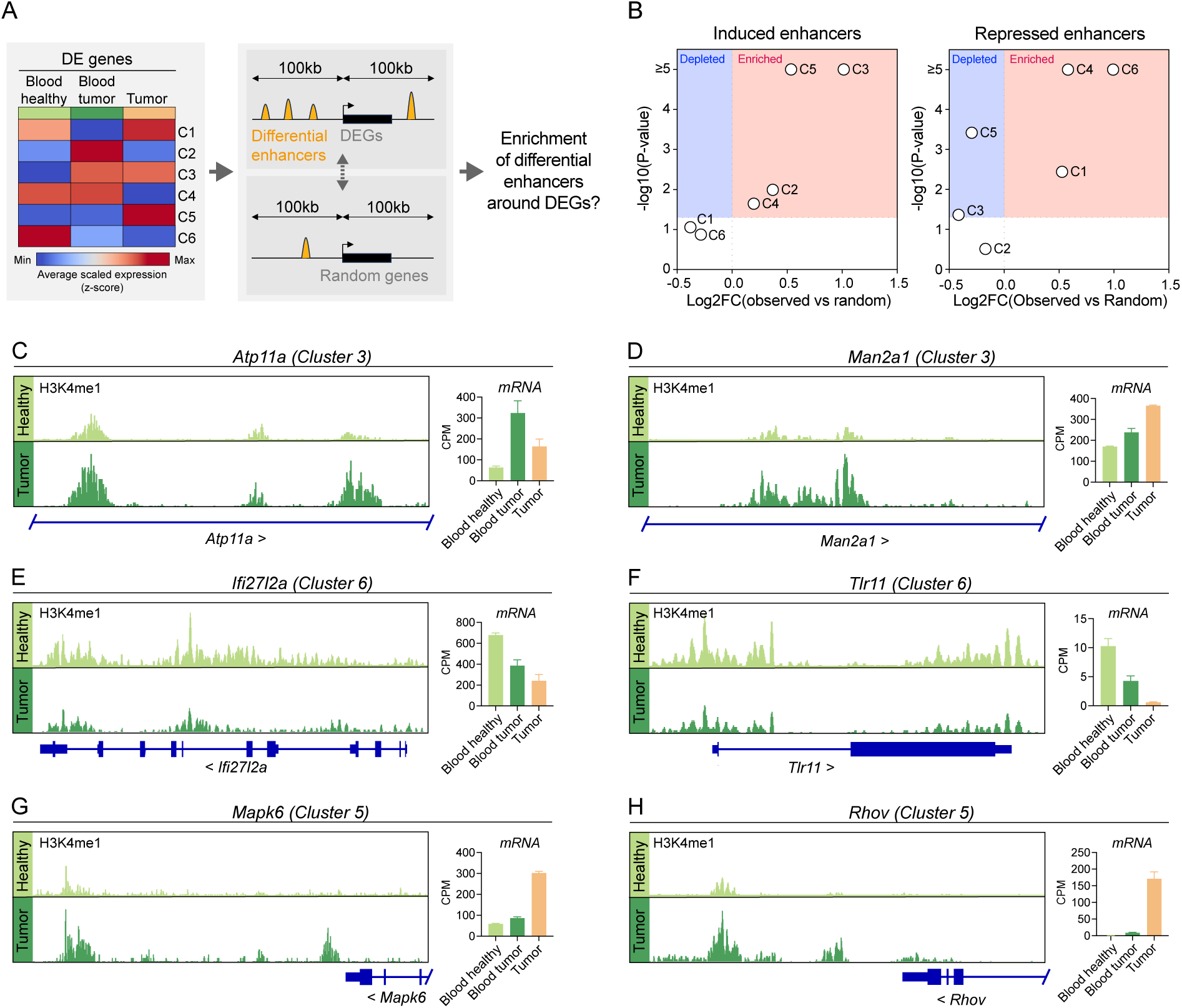
Altered enhancers in peripheral monocytes are associated with distinct gene expression patterns upon tumor infiltration. (A) Scheme summarizing the approach to determine enrichment of differential enhancers (increased or decreased H3K4me1 signal in tumor-bearing mice) +/-100 kilobase (kb) around the transcription start site of differentially expressed genes (DEGs) identified in Figure 1C. (B) Fold change and z-test p-value of the comparison of the number of differential enhancers found +/-100kb around genes belonging to the indicated expression clusters versus random gene sets of the same sizes. The number of differential enhancers found for each cluster and the corresponding random gene sets are shown in Figure S3. (C-H) Genome browser snapshots showing average H3K4me1 signal in Ly6C^hi^ monocytes from the blood of healthy and LLC tumor-bearing mice at representative genes from different expression clusters. Bar plots show mRNA expression of the same genes determined by RNA-seq. Bar plots show mean+SEM.

Overall, these results indicate that altered enhancers are enriched primarily around genes with peripheral gene expression changes that persist throughout the blood-to-tumor transition of monocytes. This suggests that epigenomic alterations can instruct sustained gene expression changes which are less likely to be reversed in the tumor microenvironment. In addition, strong enrichment of enhancers induced in the periphery around genes activated upon tumor infiltration suggest an epigenetic priming effect at these loci.

## Discussion

This study provides proof-of-principle evidence in mice for tumor-induced peripheral epigenomic alterations in monocytes and their association with gene expression changes occurring before and after tumor infiltration. Specifically, our results suggest a model whereby epigenomic programming in the periphery can shape the monocyte transcriptome in multiple ways (**Figure 5**). First, activation of promoters and/or enhancers in peripheral monocytes can initiate transcription in the periphery and maintain gene expression following tumor infiltration. Second, repression of promoters and/or enhancers in peripheral monocytes can lead to durable repression of gene expression, even after the cells infiltrate the tumor. Third, activation of certain promoters and/or enhancers in peripheral monocytes can prime genes for activation in the tumor without causing major changes in gene expression in the periphery. The latter is likely due to the fact that transcriptional activation of these genes requires enhancer binding by transcription factors that are only activated in the tumor microenvironment.

**Figure 5.**
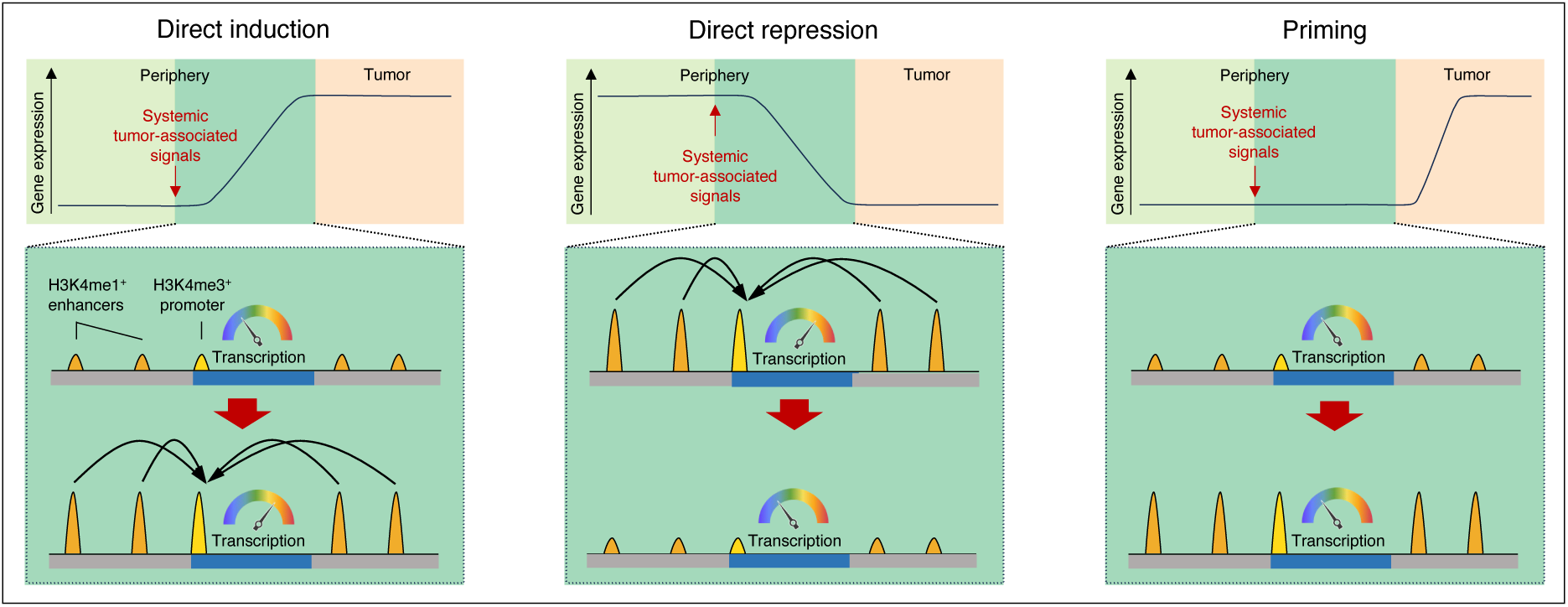
Graphical summary showing three major mechanisms by which tumor-induced epigenomic changes in peripheral monocytes can alter gene expression. Genes with direct induction show activation of promoter and/or nearby enhancers in the periphery. This is associated with increased transcription in the periphery which persists upon tumor infiltration. Genes with direct repression show the opposite pattern: repressed promoter and/or nearby enhancers are associated with durable repression of gene transcription that persist following tumor infiltration. Peripheral epigenetic priming activates promoter and/or enhancers without causing major changes in transcription of the gene. This primes the gene for expression upon tumor infiltration.

We observed that peripheral epigenomic programming of monocytes by the distant tumor involves repression of interferon-responsive promoters and enhancers harboring IRF and ISRE motifs. These findings are in line with previous observations of a suppressed interferon response of peripheral monocytes in cancer patients (16–18, 47). Our results suggest that epigenetic repression of interferon-responsive promoters and enhancers could represent one possible mechanism by which the systemic influence of a tumor desensitizes monocytes to immunostimulatory interferons and thereby suppresses anti-tumor immunity.

By integrating data on peripheral epigenomic changes and the transcriptional response during blood-to-tumor transition of monocytes, we found that genes activated in the tumor microenvironment associate with enhancers that are already induced in the periphery in response to the tumor. Generally, establishment of the enhancer repertoire is governed by lineage-specific differentiation programs and prior stimuli received by the cell (21). In macrophages, pre-established enhancers are the dominant genomic sites where signal-dependent transcription factors, such as NF-κB, AP-1 and STAT members, can bind the DNA and activate transcription in response to microenvironmental stimuli (21). Our results indicate that new H3K4me1^+^ enhancers harboring binding motifs for signal-dependent transcription factors, including AP-1 and STAT, are established in peripheral monocytes in response to the distal tumor. An important consequence of this may be an amplified transcriptional response to signals that activate these transcription factors upon tissue infiltration. We observed that the expression levels of several key transcription factors (e.g. C/EBPβ, AP-1, RUNX3) increase following tumor infiltration, which could further facilitate the activation of pre-established enhancers containing their binding motifs. It will be important to investigate to what extent such remodeling of the enhancer landscape prior to tumor infiltration can bias monocyte fate and macrophage polarity in the tumor through altering the transcriptional response to the microenvironment.

This study adds support to the emerging notion that systemic immune perturbations induced by tumor-associated signals can influence localized responses to the tumor (48). Specifically, our work uncovers epigenomic programming of monocytes as a previously unexplored tumor-induced systemic immune perturbation. Identifying the systemic tumor-associated signals driving this process will be highly relevant in the future. Plasma proteomics and metabolomics analyses to detect tumor-associated signals, followed by *in vivo* perturbations of candidate pathways combined with epigenomic and transcriptomic profiling of monocytes will likely yield valuable insights to answer this question. Tumor-derived factors reaching the systemic circulation are often dictated by the specific genetic alterations of a tumor (49). Hence, it will be interesting to determine to what extent epigenomic programming and its drivers are specific for each tumor type. This work also paves the way for future studies which should assess epigenomic alterations in peripheral monocytes in cohorts of cancer patients. Such studies could potentially identify novel epigenetic biomarkers for diagnosis and disease follow-up.

Overall, our results pinpoint the epigenomic landscape as an additional regulatory layer disrupted in monocytes by the systemic effects of a tumor. Given the importance of the epigenomic profile in controlling cell fate and responses to the microenvironment, our work has important implications for understanding how monocytes are co-opted to support cancer progression.

## Materials and Methods

### Mice

All experiments were performed with age-matched female mice. C57BL/6J mice were from Janvier. All procedures followed the guidelines of the Belgian Council for Laboratory Animal Science and were approved by the Ethical Committee for Animal Experiments of the Vrije Universiteit Brussel (licenses 15-220-3 and 19-220-8).

### Cell lines and tumor models

LLC cells were from ATCC. The cells were maintained in DMEM (Gibco) supplemented with 10% (v/v) heat-inactivated fetal calf serum (FCS; Capricorn Scientific), 300 μg/ml L-glutamine (Sigma), 100 units/ml penicillin and 100 μg/ml streptomycin (Gibco). For tumor implantation, 3×10^6^ LLC were injected subcutaneously into the right flank of mice in HBSS. Blood and tumor were collected 16-17 days after tumor implantation.

### Blood collection and tumor dissociation

Blood was collected from mice in 1 ml syringes containing 0.5 mol/L EDTA. Red blood cells were lysed using Ammonium–Chloride–Potassium (ACK) lysis buffer. Tumors were excised, cut in small pieces, incubated with 10 U/ml collagenase I (Worthington), 400 U/ml collagenase IV (Worthington) and 30 U/ml DNase I (Worthington) in RPMI for 30 min at 37°C, squashed and filtered. Following red blood cell lysis with ACK buffer, LymphoPrep density gradient was used to remove debris and dead cells.

### Cell sorting

Blood and tumor cell suspensions were resuspended in HBSS with 2 mmol/L EDTA and 0.5% (v/v) FCS. To prevent nonspecific antibody binding to Fcγ receptors, cells were pre-incubated with CD16/CD32-specific antibody (clone 2.4G2). Cell suspensions were then incubated with fluorescently labelled antibodies diluted in HBSS with 2 mml/L EDTA and 1% (v/v) FCS for 20 min at 4°C and then washed with the same buffer. The following fluorochrome-conjugated antibody clones were used: CD11b (M1/70), Ly6G (1A8), Ly6C (HK1.4), MHC-II (M5/114.15.2). Ly6C^hi^ monocytes were gated as CD11b^+^Ly6G^-^Ly6C^hi^MHC-II^-^ cells. All macrophages in LLC tumors are Ly6C^-^ and were therefore excluded (24). Fluorescence-activated cell sorting was performed using a BD FACSAria II (BD Biosciences).

### RNA-seq

RNA from sorted cells was isolated using TRIzol reagent (Invitrogen) according to the manufacturer’s instructions. RNA-Seq library was prepared using TruSeq RNA Sample Preparation Kit (Illumina) according to the manufacturer’s protocol. Libraries were sequenced with Illumina HiScanSQ sequencer.

### RNA-seq data analysis

Raw sequencing reads were processed by the nfcore RNA-seq pipeline (v3.10.1) with default parameters (50). Briefly, the reads were trimmed with trimgalore (v0.6.7) and cutadapt (v3.4) then mapped to the *mm10* genome using STAR (v2.6.1d) (51) and quantified with Salmon (v1.9.0) (52). Genes with low counts were removed using edgeR’s filterByExpr function (*min.count = 3*) (53). Principal component analysis (PCA) was performed using PCATools R package (*removeVar = 0.1*) . EdgeR was used for normalization and differential gene expression analysis with a threshold of FDR less than 0.05 and absolute log2 fold-change greater than 0.5. K-means clustering (k = 6) and heatmaps were generated using pheatmap. We used EnrichR for functional annotation of clusters against the MSigDB Hallmark 2020 database (54).

### ATAC-seq

ATAC-seq was carried out as described earlier with minor modifications (55). Monocytes were sorted into ice-cold PBS in three biological replicates. After centrifugation, the cell pellet was resuspended in ATAC lysis buffer (10 mmol/L Tris-HCl pH7.4, 10 mmol/L NaCl, 3 mmol/L MgCl2, 0.1% IGEPAL). After centrifugation, cell nuclei were tagmented using Nextera DNA Library Preparation Kit (Illumina). After tagmentation, DNA was purified with MinElute PCR Purification Kit (QIAGEN). Tagmented DNA was amplified with Kapa Hifi Hot Start Kit (Kapa Biosystems) using 9 PCR cycles. Amplified libraries were purified again with MinElute PCR Purification Kit. Fragment distribution of libraries was assessed with Agilent Bioanalyzer and libraries were sequenced on a HiSeq 2500 platform.

### ATAC-seq data analysis

Raw sequencing reads were processed by the nfcore ATAC-seq pipeline (v1.2.1) with default parameters. Briefly, the reads were trimmed with trimgalore (v0.6.4_dev) then mapped to the *mm10* genome using BWA (v0.7.17-r1188). Low quality reads and reads mapping to blacklisted regions were discarded. Duplicated reads were merged. We used MACS2 (56) to identify significantly enriched genomic regions using FDR 5%.

### H3K4me3 CUT&Run

The CUT&RUN protocol was performed using the Epicypher CUTANA CUT&RUN kit according to the manufacturer’s instructions with some minor modifications. Briefly, monocytes were recovered from freezing media by quickly thawing samples at 37°C. Cells were spun down at 600g for 3 minutes at room temperature. Cells were then resuspended in 105uL of room temperature wash buffer and cell viability was confirmed. Cells were then incubated with concavilin A beads for 10 minutes at room temperature. 0.5ug of antibodies for target proteins were then added and incubated overnight on nutator at 4°C. The following day, cells were washed with cell permeabilization buffer, before the addition of the pAG-MNase. Cells were incubated with the pAG-MNase for 10 minutes at room temperature. To digest and release target chromatin, 100mM Calcium Chloride was added to the reaction. After a 2-hour incubation on nutator at 4°C, the reaction was halted by the addition of a stop mastermix. Reactions were placed on a thermocycler at 37°C for 10 minutes. DNA was purified using SPRISelect beads (1.4X volume). Libraries were then prepared with Epicypher’s CUT&RUN Library Prep Kit according to the manufacturer’s protocol and then subjected to pair-end sequencing on a NextSeq2000 machine (1x50bp reads).

### H3K4me1 and H3K27ac CUT&Tag

Sorted Ly6C^hi^ monocytes were cryopreserved in ice cold cryopreservation solution consisting of 50% FBS, 40% RPMI and 10%DMSO. Cryopreserved samples were sent to Active Motif for CUT&Tag in two replicates. Briefly, cells were incubated overnight with Concanavalin A beads and 1 µl of the primary anti-H3K4me1 antibody per reaction (Active Motif, catalog number 39297) and 1 µl of the primary anti-H3K27ac antibody per reaction (Active Motif, catalog number 39135). After incubation with the secondary anti-rabbit antibody (1:100), cells were washed and tagmentation was performed at 37℃ using protein-A-Tn5. Tagmentation was halted by the addition of EDTA, SDS and proteinase K at 55℃, after which DNA extraction and ethanol purification was performed, followed by PCR amplification and barcoding (see Active Motif CUT&Tag kit, catalog number 53160 for recommended conditions and indexes). Following SPRI bead cleanup (Beckman Coulter), the resulting DNA libraries were quantified and sequenced on Illumina’s NextSeq 550 (PE38).

### CUT&Run and CUT&Tag data analysis

Raw sequencing reads were processed by the nfcore cutandrun pipeline (v3.0.0) with default parameters. Briefly, the reads were trimmed with trimgalore (v0.6.6) then mapped to the *mm10* genome using Bowtie2 (v2.4.4). Low quality reads and reads mapping to blacklisted regions were discarded. Duplicated reads were merged. We used MACS2 to identify significantly enriched genomic regions using FDR 5%. Genomic annotation of the peaks was done using ChIPseeker (57). We used DiffBind (filter = 3, minOverlap = 2) to identify differential genomic regions using edgeR. Distance to nearest DEG’s transcription start site (TSS) was calculated using plyranges (+/- 100kb). Coverage files were created using deepTools; normalized with RPKM. Read density plots and corresponding boxplot quantification was calculated by deepTools and profileplyr. For de novo motif enrichment, we used HOMER findMotifsGenome.pl with default parameters.

### Data availability

RNA-seq, ATAC-seq, CUT&Run and CUT&Tag datasets generated in this study will be available following peer-review at the NCBI GEO under accession number GSE268992.

## Acknowledgements

The authors thank Mikael J. Pittet and Aleksandar Murgaski for critical input on the manuscript.

## Author contributions

M. K. conceptualized and initiated the study and wrote the manuscript. M.K. and L. H. prepared the figures. M.K., D.L., L.N. and J. A.V.G. designed experiments. M.K., E.H., W.K.B., P.T. and S.P. performed experiments. A.D., J.B. and Y.E. provided technical assistance. L.H., D.K., L.M. and Y.S. performed computational analyses. B.D. and Z.C. provided critical input for research design, computational analyses and data interpretation. D.L., L.N. and J.A.V.G. obtained funding, supervised experiments and data analysis, and edited the manuscript.

---

**Figure S1.**
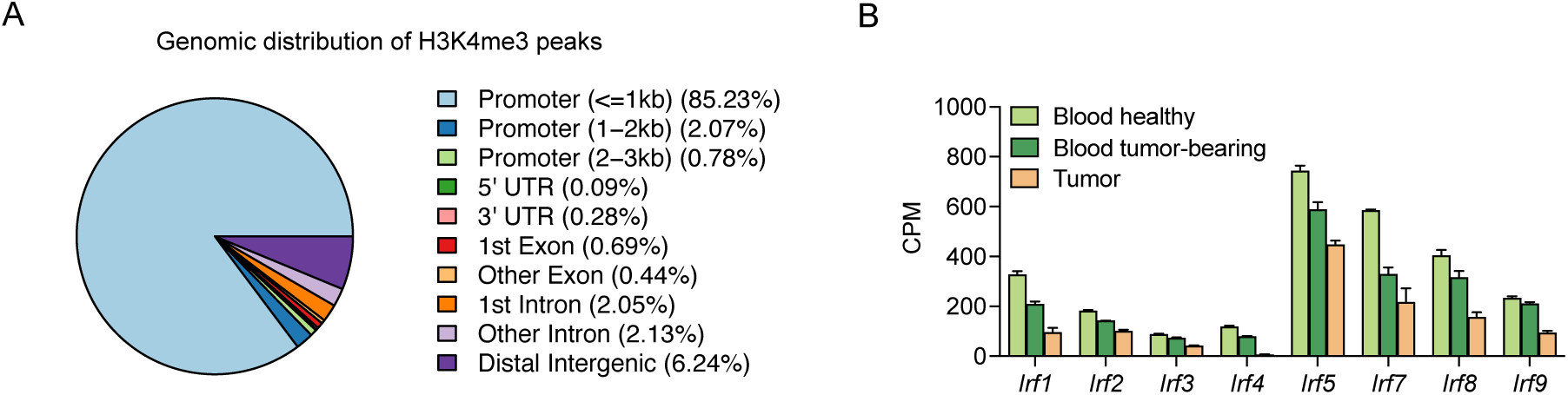
(A) Genomic distribution of H3K4me3 signal. (B) Bar graph showing expression of IRF transcription family members. Bar graph shows mean+SEM.

**Figure S2.**
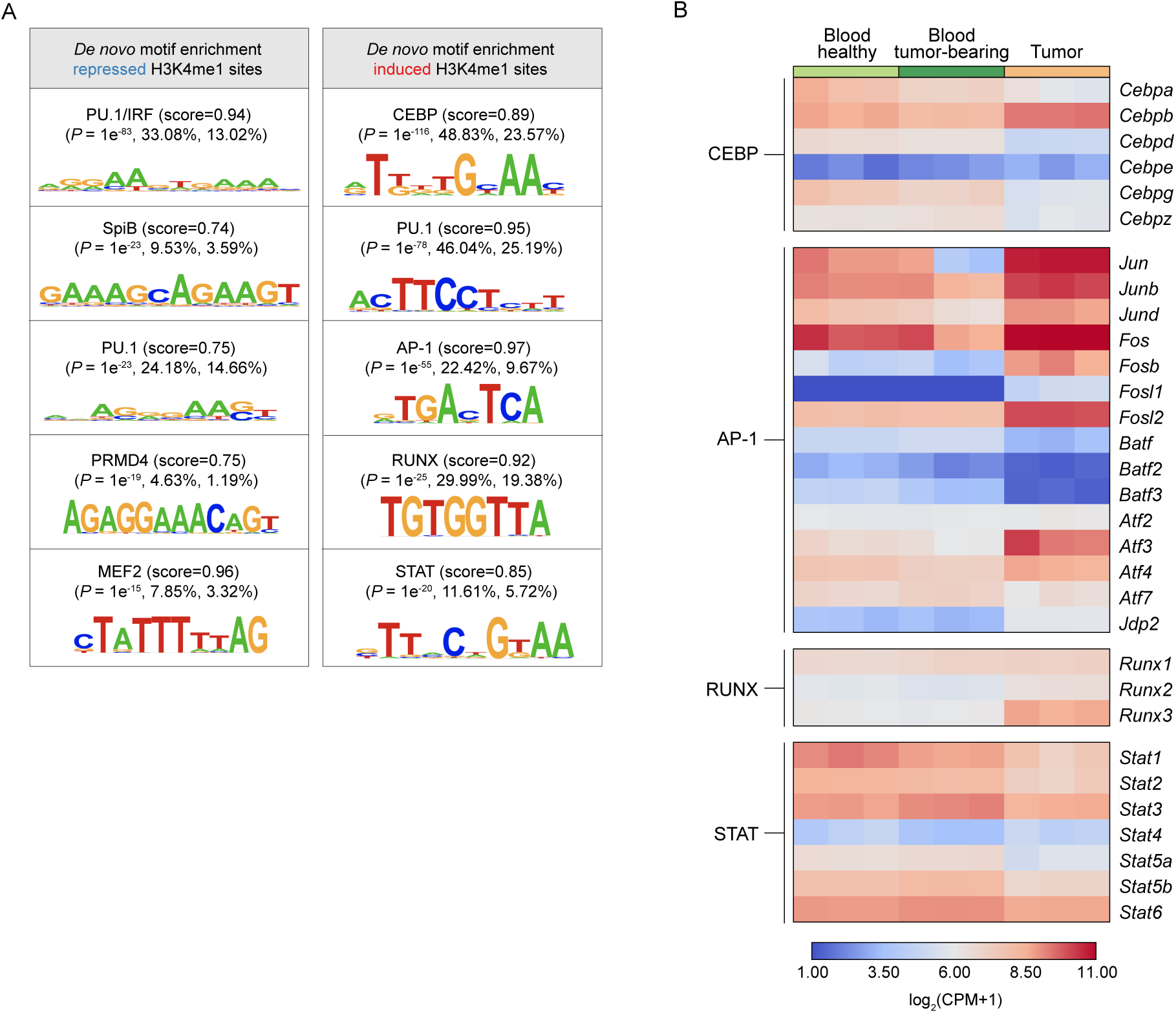
(A) Enriched *de novo* motifs among genomic sites with induced or repressed H3K4me1 (related to Figure 3E). (B) Heatmap showing expression levels of transcription factors belonging to CEBP, AP-1, RUNX and STAT families.

**Figure S3.**
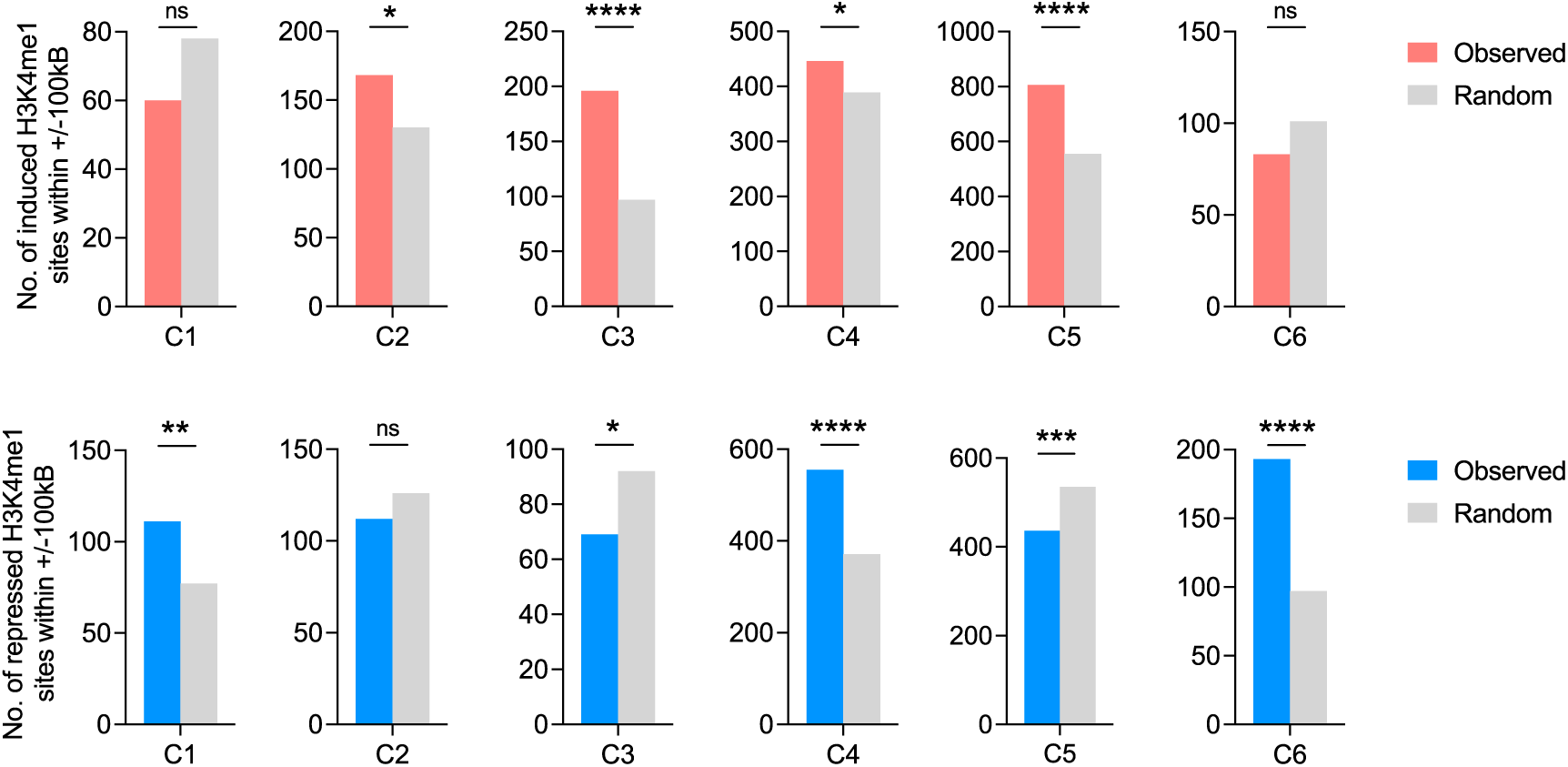
Number of enhancers with induced or repressed H3K4me1 found near (+/-100kb from the transcription start site) the different gene clusters or near random gene sets of the same size (related to Figure 4B). Z-test, ns: not significant, *P<0.05, **P<0.01, ***P<0.001.

## List of Supplementary Materials

**Table S1:** List of differentially expressed genes and their cluster identities (related to Figure 1)

**Table S2:** List of differential H3K4me3 peaks (related to Figure 2)

**Table S3:** List of differential H3K4me1 peaks (related to Figure 3)

## References

1. M. Guilliams, A. Mildner, S. Yona, Developmental and Functional Heterogeneity of Monocytes. Immunity 49, 595–613 (2018).

2. S. Singhal, et al., Human tumor-associated monocytes/macrophages and their regulation of T cell responses in early-stage lung cancer. Science Translational Medicine 11, eaat1500-16 (2019).

3. C. E. Olingy, H. Q. Dinh, C. C. Hedrick, Monocyte heterogeneity and functions in cancer. J. Leukoc. Biol. 106, 309–322 (2019).

4. R. Zilionis, et al., Single-Cell Transcriptomics of Human and Mouse Lung Cancers Reveals Conserved Myeloid Populations across Individuals and Species. Immunity 50, 1317–1334.e10 (2019).

5. E. Azizi, et al., Single-Cell Map of Diverse Immune Phenotypes in the Breast Tumor Microenvironment. Cell 1–53 (2018).

6. S. Cheng, et al., A pan-cancer single-cell transcriptional atlas of tumor infiltrating myeloid cells. Cell 184, 792–809.e23 (2021).

7. T. Kitamura, et al., Monocytes Differentiate to Immune Suppressive Precursors of Metastasis-Associated Macrophages in Mouse Models of Metastatic Breast Cancer. Frontiers in Immunology 8, 69–14 (2018).

8. S. Tadepalli, et al., Rapid recruitment and IFN-I–mediated activation of monocytes dictate focal radiotherapy efficacy. Sci. Immunol. 8, eadd7446 (2023).

9. D. Kwart, et al., Cancer cell-derived type I interferons instruct tumor monocyte polarization. Cell Rep. 41, 111769 (2022).

10. C. Krieg, et al., High-dimensional single-cell analysis predicts response to anti-PD-1 immunotherapy. Nat Med 24, 144–153 (2018).

11. T. M. Carroll, et al., Tumor monocyte content predicts immunochemotherapy outcomes in esophageal adenocarcinoma. Cancer Cell 41, 1222–1241.e7 (2023).

12. R. M. Zemek, et al., Temporally restricted activation of IFNβ signaling underlies response to immune checkpoint therapy in mice. Nat Commun 13, 4895 (2022).

13. G. Kroemer, J. L. McQuade, M. Merad, F. André, L. Zitvogel, Bodywide ecological interventions on cancer. Nat Med 29, 59–74 (2023).

14. M. Kiss, A. A. Caro, G. Raes, D. Laoui, Systemic Reprogramming of Monocytes in Cancer. Frontiers in oncology 10, 1399 (2020).

15. A. Robinson, et al., Systemic Influences of Mammary Cancer on Monocytes in Mice. Cancers 14, 833 (2022).

16. Y. Lin, et al., Immunosuppressive CD14+HLA-DRlow/− monocytes in B-cell non-Hodgkin lymphoma. Blood 117, 872–881 (2011).

17. L. Wang, et al., Breast cancer induces systemic immune changes on cytokine signaling in peripheral blood monocytes and lymphocytes. EBioMedicine 52, 102631 (2020).

18. R. N. Ramos, et al., CD163 +tumor-associated macrophage accumulation in breast cancer patients reflects both local differentiation signals and systemic skewing of monocytes. Clinical & Translational Immunology 9, 891–18 (2020).

19. L. Cassetta, et al., Human Tumor-Associated Macrophage and Monocyte Transcriptional Landscapes Reveal Cancer-Specific Reprogramming, Biomarkers, and Therapeutic Targets. Cancer Cell 1–39 (2019).

20. A. Hamm, et al., Tumour-educated circulating monocytes are powerful candidate biomarkers for diagnosis and disease follow-up of colorectal cancer. Gut 65, 990–1000 (2016).

21. C. K. Glass, G. Natoli, Molecular control of activation and priming in macrophages. Nat Immunol 17, 26–33 (2016).

22. L. B. Ivashkiv, Epigenetic regulation of macrophage polarization and function. Trends Immunol. 34, 216–223 (2013).

23. Z. Czimmerer, L. Nagy, Epigenomic regulation of macrophage polarization: Where do the nuclear receptors belong? Immunol. Rev. 317, 152–165 (2023).

24. M. Kiss, et al., IL1β Promotes Immune Suppression in the Tumor Microenvironment Independent of the Inflammasome and Gasdermin D. Cancer Immunol Res 9, 309–323 (2021).

25. F. M. Consonni, et al., Heme catabolism by tumor-associated macrophages controls metastasis formation. Nat. Immunol. 22, 595–606 (2021).

26. 26. E. Alaluf, et al., Heme oxygenase-1 orchestrates the immunosuppressive program of tumor-associated macrophages. *JCI Insight* 5 (2020).

27. D. J. Schaer, et al., Hemorrhage-activated NRF2 in tumor-associated macrophages drives cancer growth, invasion, and immunotherapy resistance. J. Clin. Investig. 134, e174528 (2024).

28. A. J. Ruthenburg, C. D. Allis, J. Wysocka, Methylation of Lysine 4 on Histone H3: Intricacy of Writing and Reading a Single Epigenetic Mark. Mol. Cell 25, 15–30 (2007).

29. N. D. Heintzman, et al., Histone modifications at human enhancers reflect global cell-type-specific gene expression. Nature 459, 108–112 (2009).

30. S. H. C. Duttke, et al., Human Promoters Are Intrinsically Directional. Mol. Cell 57, 674–684 (2015).

31. F. X. Chen, E. R. Smith, A. Shilatifard, Born to run: control of transcription elongation by RNA polymerase II. Nat. Rev. Mol. Cell Biol. 19, 464–478 (2018).

32. E. Calo, J. Wysocka, Modification of Enhancer Chromatin: What, How, and Why? Mol. Cell 49, 825–837 (2013).

33. T. A. Karakasheva, et al., CD38+ M-MDSC expansion characterizes a subset of advanced colorectal cancer patients. JCI Insight 3, 99–9 (2018).

34. T. A. Karakasheva, et al., CD38-Expressing Myeloid-Derived Suppressor Cells Promote Tumor Growth in a Murine Model of Esophageal Cancer. Cancer Research 75, 4074–4085 (2015).

35. L. Chen, et al., CD38-Mediated Immunosuppression as a Mechanism of Tumor Cell Escape from PD-1/PD-L1 Blockade. Cancer Discovery 8, 1156–1175 (2018).

36. M. D. Park, et al., TREM2 macrophages drive NK cell paucity and dysfunction in lung cancer. Nat Immunol 1–10 (2023).

37. M. Molgora, et al., TREM2 Modulation Remodels the Tumor Myeloid Landscape Enhancing Anti-PD-1 Immunotherapy. Cell 1–33 (2020).

38. Y. Katzenelenbogen, et al., Coupled scRNA-Seq and Intracellular Protein Activity Reveal an Immunosuppressive Role of TREM2 in Cancer. Cell 182, 872–885.e19 (2020).

39. R. Sun, et al., TREM2 inhibition triggers antitumor cell activity of myeloid cells in glioblastoma. Sci. Adv. 9, eade3559 (2023).

40. M. Binnewies, et al., Targeting TREM2 on tumor-associated macrophages enhances immunotherapy. Cell Rep. 37, 109844 (2021).

41. G. Wang, et al., Cutting Edge: Slamf8 Is a Negative Regulator of Nox2 Activity in Macrophages. J. Immunol. 188, 5829–5832 (2012).

42. K. Neumann, et al., Clec12a Is an Inhibitory Receptor for Uric Acid Crystals that Regulates Inflammation in Response to Cell Death. Immunity 40, 389–399 (2014).

43. M. Omatsu, et al., THBS1-producing tumor-infiltrating monocyte-like cells contribute to immunosuppression and metastasis in colorectal cancer. Nat. Commun. 14, 5534 (2023).

44. O. M. Pello, et al., In Vivo Inhibition of c-MYC in Myeloid Cells Impairs Tumor-Associated Macrophage Maturation and Pro-Tumoral Activities. PLoS ONE 7, e45399 (2012).

45. O. M. Pello, et al., Role of c-MYC in alternative activation of human macrophages and tumor-associated macrophage biology. Blood 119, 411–421 (2012).

46. J. C. Black, C. Van Rechem, J. R. Whetstine, Histone Lysine Methylation Dynamics: Establishment, Regulation, and Biological Impact. Mol. Cell 48, 491–507 (2012).

47. E. Verronèse, et al., Immune cell dysfunctions in breast cancer patients detected through whole blood multi-parametric flow cytometry assay. Oncoimmunology 5, e1100791 (2015).

48. K. J. Hiam-Galvez, B. M. Allen, M. H. Spitzer, Systemic immunity in cancer. Nat. Rev. Cancer 21, 345–359 (2021).

49. M. D. Wellenstein, K. E. de Visser, Cancer-Cell-Intrinsic Mechanisms Shaping the Tumor Immune Landscape. Immunity 48, 399–416 (2018).

50. P. A. Ewels, et al., The nf-core framework for community-curated bioinformatics pipelines. Nat. Biotechnol. 38, 276–278 (2020).

51. A. Dobin, et al., STAR: ultrafast universal RNA-seq aligner. Bioinformatics 29, 15–21 (2013).

52. R. Patro, G. Duggal, M. I. Love, R. A. Irizarry, C. Kingsford, Salmon provides fast and bias-aware quantification of transcript expression. Nat. Methods 14, 417–419 (2017).

53. M. D. Robinson, D. J. McCarthy, G. K. Smyth, edgeR: a Bioconductor package for differential expression analysis of digital gene expression data. Bioinformatics 26, 139–140 (2010).

54. M. V. Kuleshov, et al., Enrichr: a comprehensive gene set enrichment analysis web server 2016 update. Nucleic Acids Res. 44, W90–W97 (2016).

55. J. D. Buenrostro, P. G. Giresi, L. C. Zaba, H. Y. Chang, W. J. Greenleaf, Transposition of native chromatin for fast and sensitive epigenomic profiling of open chromatin, DNA-binding proteins and nucleosome position. Nature Methods 10, 1213–1218 (2013).

56. Y. Zhang, et al., Model-based analysis of ChIP-Seq (MACS). Genome Biol. 9, R137 (2008).

57. Q. Wang, et al., Exploring Epigenomic Datasets by ChIPseeker. Curr. Protoc. 2, e585 (2022).

